# Binocular visual experience and sleep promote visual cortex plasticity and restore binocular vision in a mouse model of amblyopia

**DOI:** 10.1101/2022.01.25.477697

**Authors:** Jessy D. Martinez, Marcus J. Donnelly, Donald S. Popke, Daniel Torres, Lydia G. Wilson, William P. Brancaleone, Brittany C. Clawson, Sha Jiang, Sara J. Aton

## Abstract

Amblyopia arises from an altered balance of input from the two eyes to the binocular zone of primary visual cortex (bV1) during childhood, causing long-lasting visual impairment. Amblyopia is commonly treated by patching the dominant eye, however, the relative impacts of monocular vs. binocular visual experiences on restoration of bV1 function remains unclear. Moreover, while sleep has been implicated in V1 plasticity in response to vision loss, its role in recovery from amblyopia is unknown. We used monocular deprivation (MD) in juvenile mice to model amblyopia in bV1. We compared recovery of visual responses for the two eyes among bV1 regular spiking (RS, putative principal) neurons and fast-spiking (FS, putative parvalbumin-expressing [PV+]) interneurons after identical-duration, identical-quality binocular recovery (BR) or monocular, reverse occlusion (RO) experiences. We find that BR is quantitatively superior to RO with respect to renormalizing both bV1 populations’ visual responses. However, this recovery was seen only in freely-sleeping mice; post-BR sleep deprivation prevented functional recovery. Thus, both binocular visual experience and subsequent sleep are required to optimally renormalize bV1 responses in a mouse model of amblyopia.

## Introduction

Experience-driven synaptic plasticity during critical developmental periods affects lifelong sensory and behavioral functions (1, 2). Brief monocular deprivation (MD; occlusion of one of the two eyes) during early postnatal development shifts responsiveness of primary visual cortex (V1) neurons to lose binocular responsiveness (3, 4). This phenomenon, known as ocular dominance plasticity (ODP), results from depression of deprived eye (DE) responses, followed by potentiation of spared eye (SE) responses, among V1 neurons (5, 6). These changes are associated with a transient decrease in cortical inhibition, initiated by MD itself (7, 8). Closure of the critical period for ODP is thought to involve restoration of “mature” levels of cortical inhibition, which disrupts subsequent competitive plasticity of excitatory inputs (9-12).

ODP is a well-established model of amblyopia, a visual disorder which is caused by imbalanced input to V1 from the two eyes in childhood. Amblyopia leads to long-term disruption of binocular vision and poor visual acuity in adulthood (13-16). Standard clinical interventions to promote vision recovery in amblyopia include dominant eye patching and – more recently - intensive binocular experience (17-22). There remains significant debate regarding whether binocular or monocular interventions are superior at restoring vision to amblyopic children – with direct comparisons in patient populations in randomized clinical trials yielding conflicting results (14, 22, 23). Animal models have also been used to address this question, to assess how visual experiences affect recovery of V1 function following MD. For example, reverse occlusion (RO; analogous to dominant eye patching) induces slower (or less) recovery in developing cat V1 than binocular recovery (BR) after a period of MD, suggesting that temporally correlated input to the two eyes benefits overall recovery (24-28). Moreover, a single day of BR has been shown to restore binocularity of intrinsic signal responses in mouse V1 after MD, with a day of RO has less effect (29). Available electrophysiological data on visual recovery in adult rodents following MD suggests a similar benefit to BR over RO (30-32). However, it is unclear what changes to the V1 network (e.g. in the visual responses of excitatory vs. inhibitory neurons) mediate these differences. It is also unclear whether during the critical period, multi-day BR and RO experiences of identical duration and quality (e.g., with identical contrast and spatial frequency) have quantitatively different effects on these neuronal responses.

Sleep has been shown to benefit processes relying on synaptic plasticity, including ODP in V1 (7, 33-38). In cat V1, initial shifts in ocular dominance following a brief period of MD are augmented by a few hours of subsequent sleep (36) and are disrupted by sleep deprivation (SD) (35). This might suggest that sleep following visual experience could play a vital role in recovery of visual function in amblyopia. However, very little is known about the role of sleep in, promoting V1 functional recovery following MD. In a single study in critical period cats, a period of sleep following a brief interval of post-MD RO actually impaired (rather than enhanced) recovery of normal V1 ocular dominance (39). Thus the function of appropriately-timed sleep in promoting (or disrupting) vision recovery in amblyopia is completely unknown.

To address these questions, we first directly compared how multi-day, post-MD BR and RO - of identical duration and visual stimulus content - affect recovery of function in mouse binocular V1 (bV1). Using single-neuron recordings, we find that bV1 ocular dominance shifts caused by 5-day MD are completely reversed by a period of visually-enriched BR experience, but are only partially reversed by RO of similar duration and quality. These differential effects were observed in both regular spiking (RS) neurons and fast spiking (FS; putative parvalbumin-expressing [PV+]) interneurons. BR, but not RO, reversed MD-induced depression of DE-driven firing rate responses in both RS neurons and FS interneurons, and increases in SE-driven responses in both RS and FS populations. Recovery of function was confirmed with immunohistochemistry for DE-driven cFos expression in bV1 PV+ interneurons. DE-driven cFos expression across layers 2/3, and particularly in PV+ interneurons, was reduced after MD, and recovered to control levels after BR, but not RO. Critically, BR-driven recovery of ocular dominance, visual response changes, and DE-driven cFos expression were all disrupted by SD in the hours immediately following periods of BR. Together these results suggest that optimal recovery of bV1 function after a period of MD requires both enriched binocular visual experience and subsequent undisturbed sleep. These data have important implications for the treatment of amblyopia, which results from similar ODP mechanisms to those present in bV1 during and after MD.

## Results

### Binocular recovery (BR) causes more complete reversal of MD-induced bV1 ocular dominance shifts more than identical-duration reverse occlusion (RO)

We first directly compared the degree of visual recovery induced by multi-day BR and RO in bV1 neurons following a 5-day period of MD (**Fig. 1A**). The duration and timing of MD (P28-33; during the peak of the critical period for ODP) was chosen with the aim of inducing a robust ocular dominance shift, with changes to both DE and SE responses in bV1 (5). To ensure comparable quality and duration of visual experience between BR and RO recovery groups, from P33-38, these mice were placed for 4 h/day (starting at lights-on) in a square chamber surrounded by four LED monitors presenting high-contrast, phase-reversing gratings (8 orientations, 0.05 cycles/deg, reversing at 1 Hz) in an interleaved manner. During this period of visual enrichment, mice had access to a running wheel, manipulanda, and treats *(31)* in order to increase wake time and promote more consistent visual stimulation. After the 5-day recovery period, we compared bV1 neurons’ visual responses for stimuli presented to either the right or left eyes, for the hemisphere contralateral to the original DE.

**Fig. 1.**
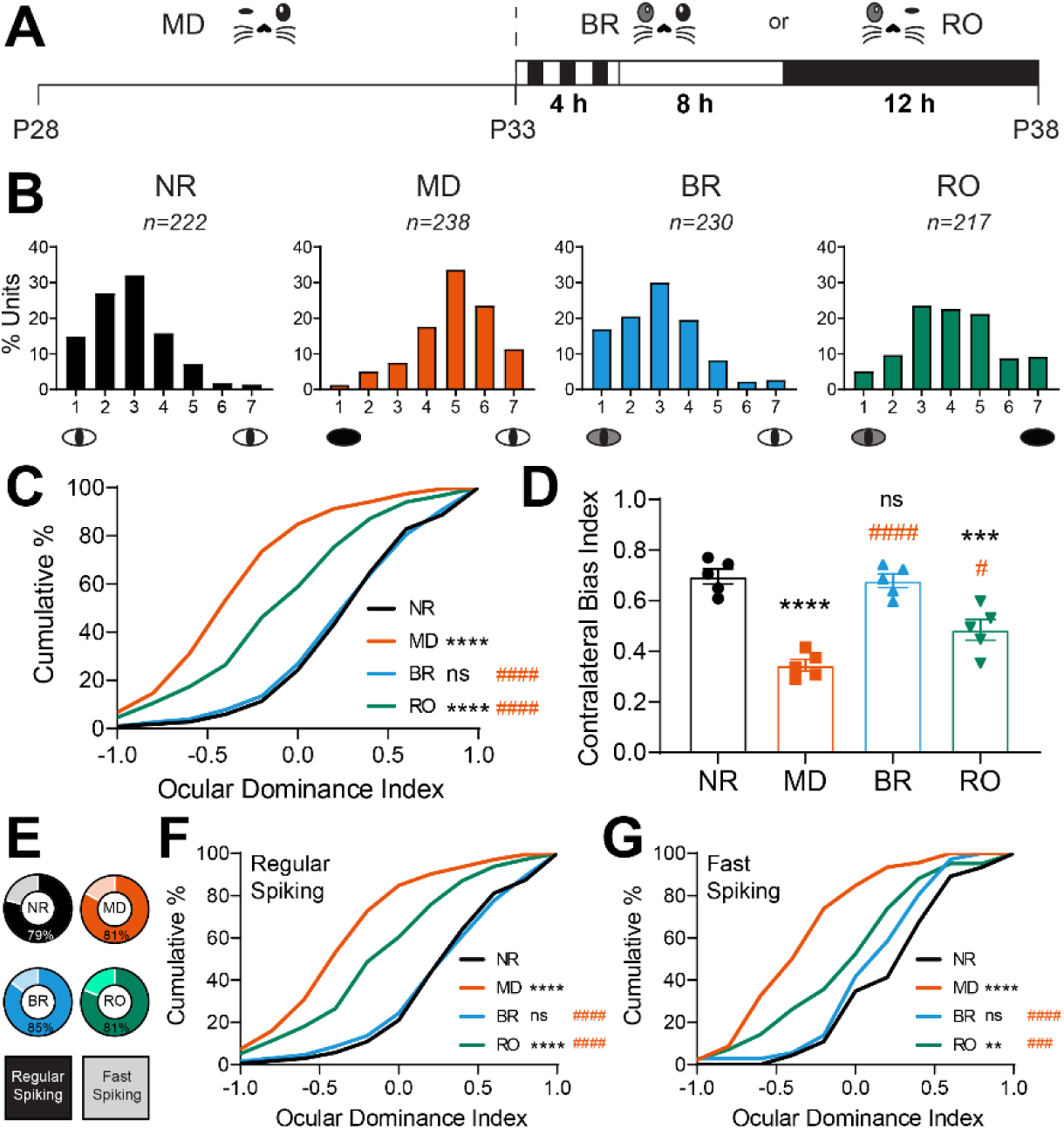
BR is more effective than RO at reversing MD-induced ocular dominance shifts. **(A)** Experimental design. Mice underwent 5-day MD from P28-P33. MD mice were recorded at P33. Two recovery groups with either binocular recovery (BR) or reverse occlusion (RO) visual experience from P33-38 had daily 4-h periods of visual enrichment starting at lights on and were recorded at P38. Normally-reared (NR) mice were recorded at P38 without prior manipulation of vision. **(B)** Ocular dominance histograms from bV1 neurons recorded contralateral to the original DE for all four groups, using a 7-point scale (1= neurons driven exclusively by contralateral eye; 7= neurons driven exclusively by ipsilateral eye, 4= neurons with binocular responses) *n* = 5 mice/group. **(C)** Cumulative distribution of ocular dominance indices for all neurons recorded in each group. **** (black) indicates *p* < 0.0001, K-S test vs. NR; #### (red) indicates *p* < 0.0001, K-S test vs MD; ns indicates not significant. **(D)** Contralateral bias indices for mice in each treatment group. One-way ANOVA: *F* (3, 16) = 29.34, *p* < 0.0001. Tukey’s *post hoc* test vs NR – MD: *p* < 0.0001; BR: *p* = ns; RO: *p* < 0.001. Tukey’s *post hoc* test vs MD – BR: *p* < 0.0001; RO: *p* < 0.01. Error bars indicate mean ± SEM. **(E)** The proportion of recorded neurons classified as regular spiking (RS) neurons and fast-spiking (FS) interneurons in each treatment group. RS neurons: NR (*n* = 175); MD (*n* = 192); BR (*n* = 196); RO (*n* = 175). FS interneurons: NR (*n* = 47); MD (*n* = 46); BR (*n* = 34); RO (*n* = 42). **(F-G)** Ocular dominance index cumulative distributions for RS neurons **(F)** and FS interneurons **(G)**. Ocular dominance index values for both populations were significantly shifted in favor of the SE after MD, were comparable to those of NR mice after BR, and were intermediate – between NR and MD values – after RO. ** and **** (black) indicate *p* < 0.01 and *p* < 0.0001, K-S test vs. NR; ### and #### (red) indicate *p* < 0.001 and *p* < 0.0001, K-S test vs MD.

Consistent with previous reports *(1, 5)*, 5-day MD induced a large ocular dominance shift in favor of the SE compared to normally-reared (NR) control mice with binocular vision from birth (**Fig. 1B-D**). 5 days of BR visual experience returned bV1 ocular dominance to a distribution similar to age-matched NR mice, completely reversing the effects of MD. After BR, ocular dominance index distributions (**Fig. 1C**) and contralateral bias indices for each mouse (**Fig. 1D**) matched those of NR mice, showing a preference for the DE (contralateral) eye. In contrast, ocular dominance distributions following 5-day RO visual experience were intermediate between MD mice and age-matched NR mice (**Fig. 1B-D**), suggesting only partial recovery.

MD is known to effect a change in the balance of activity between principal (RS; mainly glutamatergic) neurons and FS (mainly PV+, GABAergic) interneurons (7, 8, 40). In our extracellular recordings, FS interneurons (identifiable based on distinctive spike waveform features (7, 41)) represented roughly 15-20% of all stably-recorded neurons, across all treatment conditions (**Fig. 1E**). We found that relative to neurons recorded from NR mice, MD led to similar ocular dominance shifts toward the SE in both RS neurons and FS interneurons (**Fig. 1F** and **1G**). These MD-induced changes were completely reversed in both RS and FS populations in BR mice, but were only partially reversed in RO mice (**Fig. 1F** and **1G**). We conclude that with respect to ocular dominance distributions in bV1, 5-day BR is quantitatively superior to 5-day RO at reversing effects of MD.

### BR and RO differentially restore bV1 RS neuron and FS interneuron firing rate responses after MD

MD leads to sequential changes in V1 neurons’ responses to DE and SE stimulation (which are depressed and potentiated, respectively) (5-7, 36, 42). We next investigated which of these changes could be reversed in bV1 neurons as a function of post-MD BR or RO. Consistent with previous reports (5, 6), 5-day MD reduced the magnitude of DE-driven visually-evoked firing rate responses across bV1. Depression of DE responses was partially reversed after either BR or RO recovery experience (**Fig. 2 – Figure supplement 1A**). Potentiation of SE responses was partially reversed by RS, and completely reversed after BR (**Fig. 2 – Figure supplement 1B**). To better characterize microcircuit-level changes due to MD, we examined how DE and SE response recovery varied between RS neuron and FS interneuron populations, and in different layers of bV1. DE responses were significantly depressed after 5-day MD as previously reported (5, 6); these changes were seen across cortical layers, in both RS neurons (**Fig. 2A top**, and **2B**) and FS interneurons (**Fig. 2D top** and **2E**). In both populations, DE response depression was most pronounced in the extragranular layers. Both BR and RO both largely reversed DE response depression in RS neurons, although modest differences remained after RO (**Fig. 2A top**); recovery appeared most complete in RS neurons in layers 5/6 (**Fig. 2B**). DE response depression in FS interneurons was fully reversed by 5-day BR (**Fig. 2D top**), with the most dramatic changes occurring in the extragranular layers (**Fig. 2E**). In comparison, response depression reversal was more modest (though still significant) after RO (**Fig. 2D top**), with the largest changes occurring in layer 4 FS interneurons (**Fig. 2E**).

**Fig. 2.**
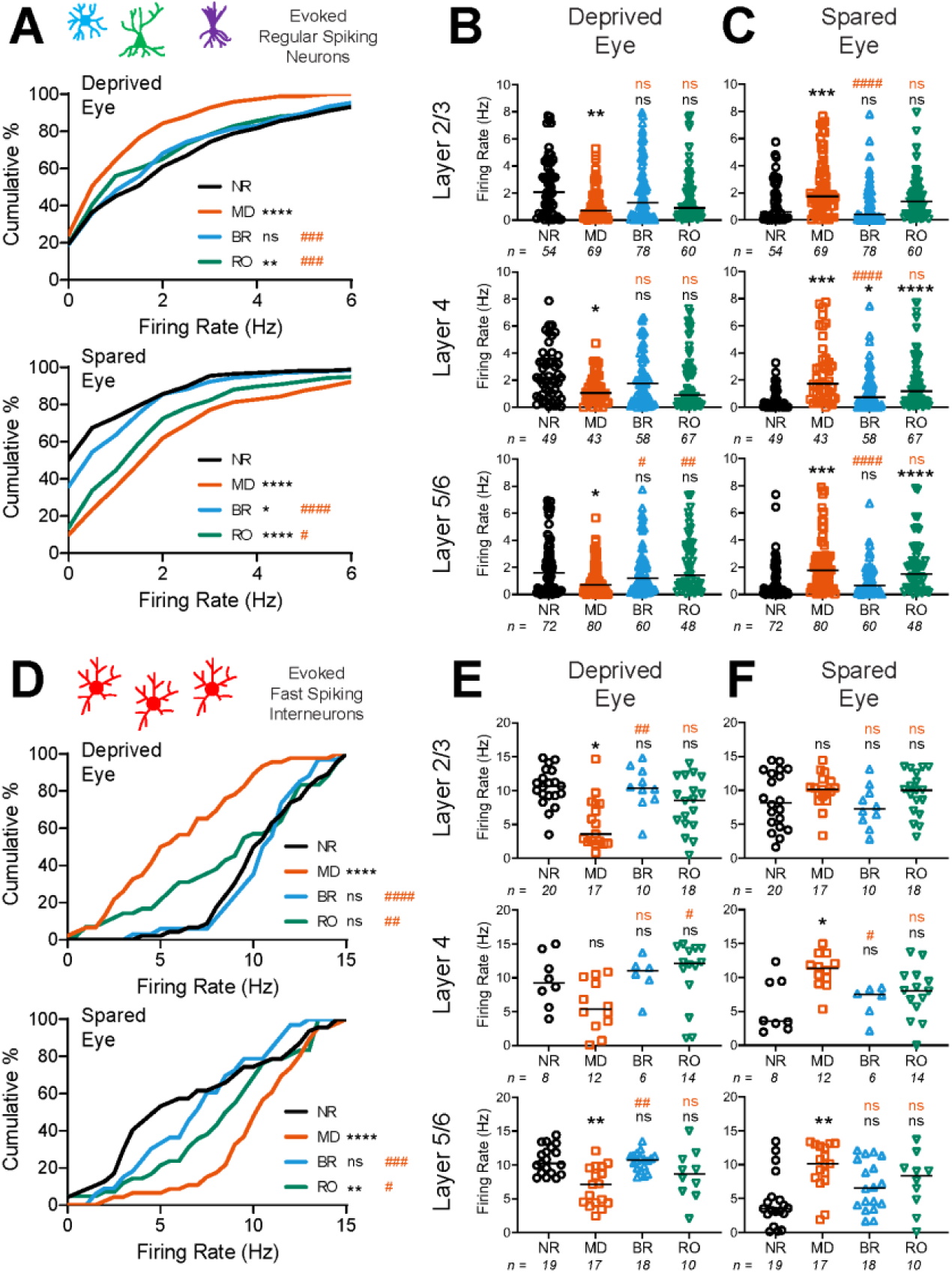
BR and RO differentially reverse MD-induced changes in DE and SE firing rate responses among RS neurons and FS interneurons. **(A)** Cumulative distributions of maximal DE **(top)** and SE **(bottom)** visually-evoked firing rate responses for bV1 RS neurons. DE responses were significantly depressed after 5-day MD; this was reversed fully after BR and partially after RO. SE responses in RS neurons showed post-MD potentiation, which was maintained after RO, but largely reversed by BR. *, **, and **** (black) indicate *p* < 0.05, *p* < 0.01, and *p* < 0.0001, K-S test vs. NR; #, ### and #### (red) indicate *p* < 0.05, *p* < 0.001, and *p* < 0.0001, K-S test vs MD. **(B-C)** RS neurons’ DE **(B)** and SE **(C)** visually-evoked responses recorded from neurons in bV1 layers 2/3, 4, or 5/6. Solid lines indicate median values for each neuron population. *, **, ***, and **** (black) indicate *p* < 0.05, *p* < 0.01, *p* < 0.001, and *p* < 0.0001, Dunn’s *post hoc* test vs. NR; #, ##, and #### (red) indicate *p* < 0.05, *p* < 0.01 and *p* < 0.0001, Dunn’s *post hoc* test vs MD. *p* < 0.01, *<* 0.05, and < 0.01 for DE responses recorded in layers 2/3, 4, and 5/6, respectively; *p* < 0.0001 for SE responses recorded in all layers, Kruskal-Wallis test. **(D)** Cumulative distributions of maximal DE **(top)** and SE **(bottom)** visually-evoked firing rate responses for FS interneurons. DE and SE responses were depressed and potentiated, respectively, after MD. These response changes were partially reversed by RO, and fully reversed by BR. ** and **** (black) indicate *p* < 0.01 and *p* < 0.0001, K-S test vs. NR; #, ##, ###, and #### (red) indicate *p* < 0.05, *p* < 0.01, *p* < 0.01, and *p* < 0.0001, K-S test vs MD. **(E-F)** FS interneurons’ DE **(E)** and SE **(F)** visually-evoked responses recorded from neurons in each bV1 layer. * and ** (black) indicate *p* < 0.05 and *p* < 0.01, Dunn’s *post hoc* test vs. NR; # and ## (red) indicate *p* < 0.05 and *p* < 0.01, Dunn’s *post hoc* test vs MD. *p* < 0.001, < 0.05, < 0.001 for DE responses recorded in layers 2/3, 4, and 5/6, respectively; *p* = ns, < 0.01, and < 0.01 for SE responses recorded in layers 2/3, 4, and 5/6 respectively, Kruskal-Wallis test.

**Fig. 2 – Figure supplement 1.**
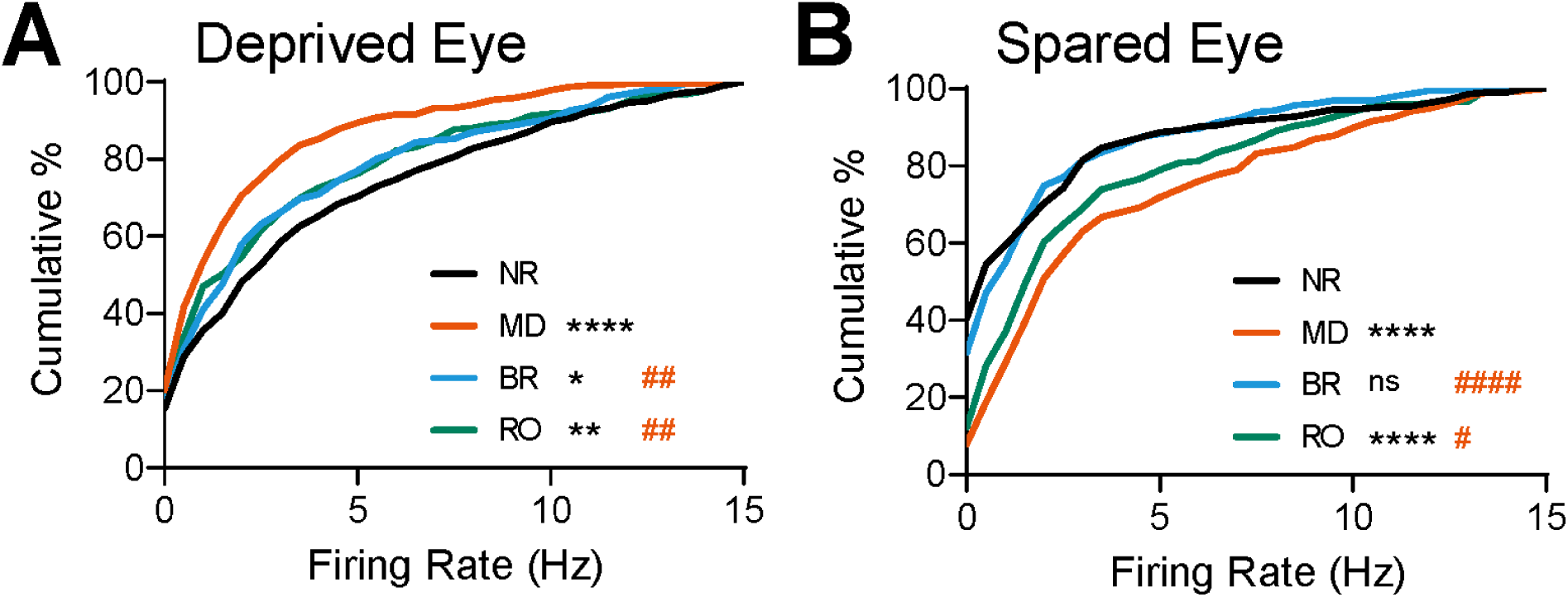
DE and SE maximal evoked firing rate distributions for all recorded bV1 neurons. Cumulative distributions of maximal DE **(A)** and SE **(B)** visually-evoked firing rate responses for RS neurons and FS interneurons combined. *, **, and **** (black) indicate *p* < 0.05, *p* < 0.01 and *p* < 0.0001, K-S test vs. NR; #, ##, and #### (red) indicates *p* < 0.05, *p* < 0.01, and *p* < 0.0001, K-S test vs MD.

MD strongly potentiated responses to SE stimulation, across both bV1 neuron populations, and across cortical layers (**Fig. 2A bottom, 2C, 2D bottom**, and **2F**). BR and RO had differential effects with respect to reversing MD-potentiated responses. For both RS neurons and FS interneurons (**Fig. 2A bottom** and **2D bottom**), potentiation of SE responses was almost completely reversed by BR. In contrast, in both neuron populations, RO led to only partial reversal of MD-induced SE response potentiation (**Fig. 2A bottom** and **2D bottom**). After BR, reversal of SE response potentiation was present in RS neurons across bV1 layers. In contrast, after RO, SE responses remained significantly potentiated in layer 4 and layers 5/6 (**Fig. 2C**). Among FS interneurons, BR tended to reverse SE response potentiation more completely than RO across all layers of bV1, with the most complete reversal (leading to significant differences from MD alone) seen in layer 4 (**Fig. 2F**).

### BR, but not RO, fully restores DE-driven cFos expression in bV1 layers 2/3

To further characterize how MD, BR, and RO affect visual responses throughout bV1, we used immunohistochemistry to quantify DE-driven cFos expression in PV+ interneurons and non-PV+ neurons. Mice were treated as shown in **Fig. 1**, after which they were returned to the visual enrichment arena for 30 min of visual stimulation of the DE only, then were perfused 90 min later. Visually-driven expression of cFos and PV expression were quantified across the layers of bV1 contralateral to the DE. Consistent with previous reports (43, 44), DE-driven cFos expression was significantly reduced across bV1 after MD (**Fig. 3A**). Both total density of cFos+ neurons and the density of cFos+ PV+ interneurons decreased after MD. BR reversed these changes, restoring DE-driven cFos expression to levels seen in NR control mice (**Fig. 3A** and **3C**). In contrast, and consistent with data shown in **Fig. 2A** and **2D**, both total DE-driven cFos expression and density of cFos+ PV+ interneurons remained significantly reduced after RO. Quantification of cFos and PV by layer showed that the largest differential effects of visual experience were seen in layers 2/3. Following MD, DE-driven cFos expression was reduced across all layers (**Fig. 3D**), and cFos+ PV+ interneuron density was dramatically reduced in layers 2/3 (and to a lesser extent, layer 4) (**Fig. 3F**). RO restored DE-driven cFos expression in layer 4 and layers 5/6, but not layers 2/3 (**Fig. 3D** and **3F**). After BR, total and PV+ interneuron cFos expression was renormalized across all layers, including layer 2/3, where cFos+ neuron density was restored to levels seen in NR mice. Together, these data suggest that activity-driven plasticity in layer 2/3, particularly in PV+ interneurons, may differ dramatically during monocular vs. binocular recovery from MD.

**Fig. 3.**
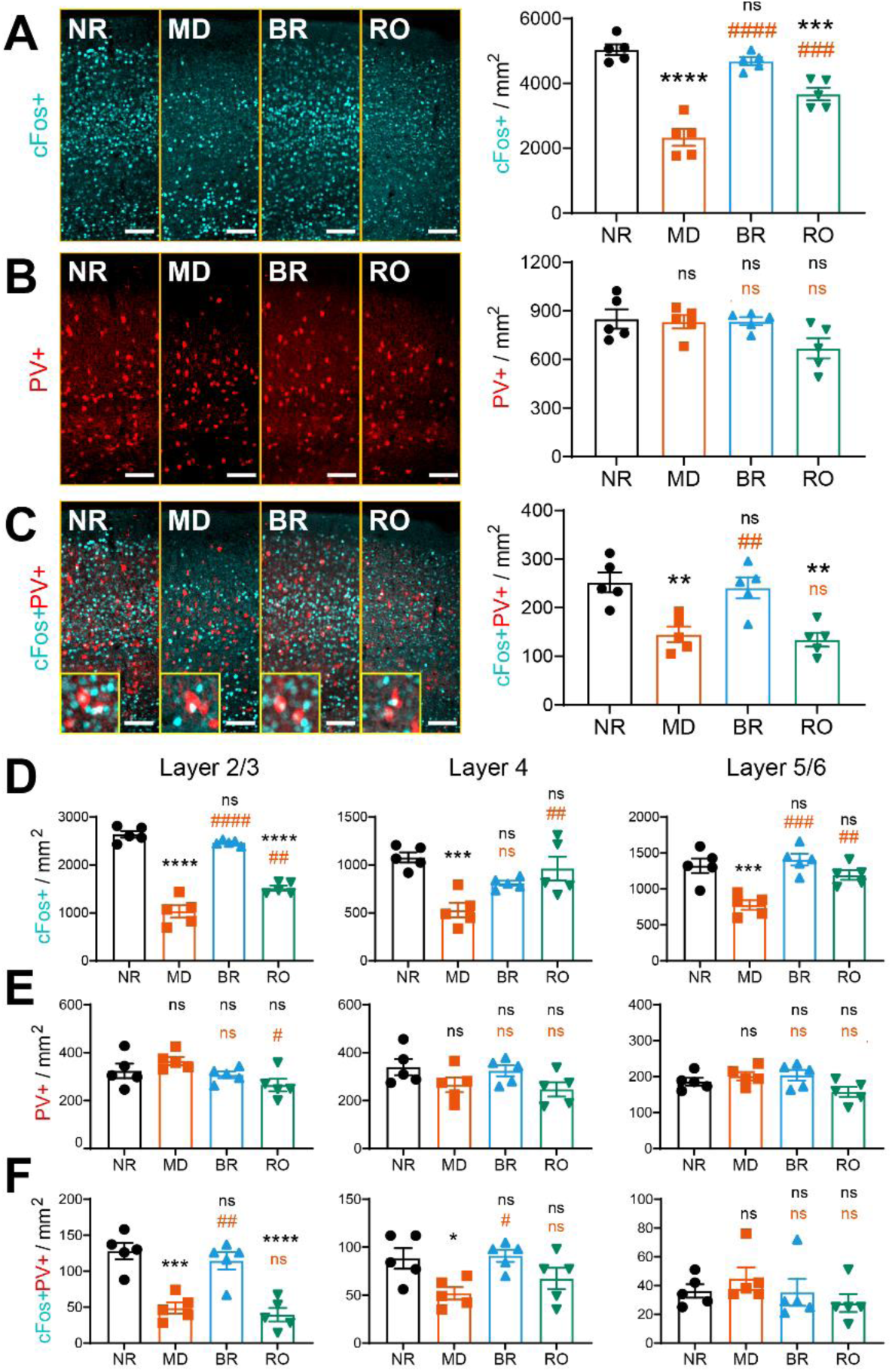
DE-driven cFos expression is reduced after MD and restored after BR, but not RO. **(A)** Representative images of bV1 cFos (cyan) across treatment groups following DE stimulation. Mice (*n* = 5/treatment group) received DE-only visual stimulation for 30 min, then were returned to their home cages for 90 min prior to perfusion. DE-driven cFos+ neuron density was decreased in bV1 after MD. cFos expression was fully rescued after BR and partially rescued after RO. One-way ANOVA: *F* (3, 16) = 39.65, *p* < 0.0001; *** and **** (black) indicate *p* < 0.001 and *p* < 0.0001, Tukey test vs. NR; ### and #### (red) indicate *p* < 0.001 and *p* < 0.0001, Tukey test vs MD. **(B)** Representative image of parvalbumin (PV) [red] following DE stimulation. Density of PV+ bV1 interneurons was similar between groups. One-way ANOVA: *F* (3, 16) = 2.99, p = 0.062. **(C)** cFos+ PV+ interneuron density decreased with MD and recovered with BR, but not RO. One-way ANOVA: *F* (3, 16) = 11.40, *p* = 0.0003; ** (black) indicates *p* < 0.01, Tukey test vs. NR; ## (red) indicates *p* < 0.01, Tukey test vs MD. **(D)** cFos+ neuron density in bV1 layers 2/3, 4, and 5/6. One-way ANOVA for layers 2/3, 4, or 5/6, respectively: *F* (3, 16) = 95.41, *p* < 0.0001, *F* (3, 16) = 9.093, *p* = 0.001, and *F* (3, 16) = 12.35, *p* = 0.0002. **(E)** PV+ interneuron density in bV1 layers 2/3, 4, and 5/6. One-way ANOVA for layers 2/3, 4, or 5/6, respectively: *F* (3, 16) = 3.40, *p* = 0.044, ns, and ns. **(F)** cFos+PV+ interneuron density in bV1 layers 2/3, 4, and 5/6. One-way ANOVA for layers 2/3, 4, or 5/6, respectively: *F* (3, 16) = 18.88, *p* < 0.0001, *F* (3, 16) = 4.25, *p* = 0.022, and ns. *, ***, and **** (black) indicate *p* < 0.05, *p* < 0.001, and *p* < 0.0001, Tukey test vs. NR; #, ##, ### and #### (red) indicate *p* < 0.05, *p* < 0.01, *p* < 0.001, and *p* < 0.0001, Tukey test vs MD. Error bars indicate mean ± SEM. Scale bar = 100 μm.

### Sleep in the hours following BR visual experience is necessary for ocular dominance recovery

Initial shifts in ocular dominance in favor of the SE are promoted by periods of sleep following monocular visual experience (7, 35, 36, 45). However, it is unclear whether, or how, sleep contributes to bV1 functional recovery after MD. Because 5-day BR (with 4 h of binocular visual enrichment per day) was effective at reversing many of the effects of prior MD, we tested whether post-visual enrichment sleep plays an essential role in this recovery. Mice underwent the same 5-day MD and 5-day BR periods shown in **Fig. 1**. Following each daily visual enrichment period, mice were returned to their home cage, and over the next 4 h were either sleep deprived (SD) under dim red light (to prevent additional visual input to V1) or allowed *ad lib* sleep (BR+SD and BR+Sleep, respectively; **Fig. 4A**). We then compared bV1 neurons’ visual responses for stimuli presented to either the right or left eyes, for the hemisphere contralateral to the original DE, between BR+Sleep and BR+SD mice. In contrast to prior reports on the effects of SD following RO in critical period cats (39), we found that SD in the hours following daily BR visual experience reduced post-MD recovery of ocular dominance in favor of the original DE (**Fig. 4B**). Ocular dominance index and contralateral bias index values for bV1 neurons recorded from BR+SD mice were significantly reduced compared to those of BR+Sleep mice, indicating reduced DE preference similar to that seen after MD alone (**Fig. 4C-D**). These effects of SD on ocular dominance recovery across BR were present in both RS neurons and FS interneurons in bV1 (**Fig. 4E-G**). Thus, in the context of BR-mediated recovery from MD, post-experience sleep plays an essential role in recovery of ocular dominance in bV1.

**Fig. 4.**
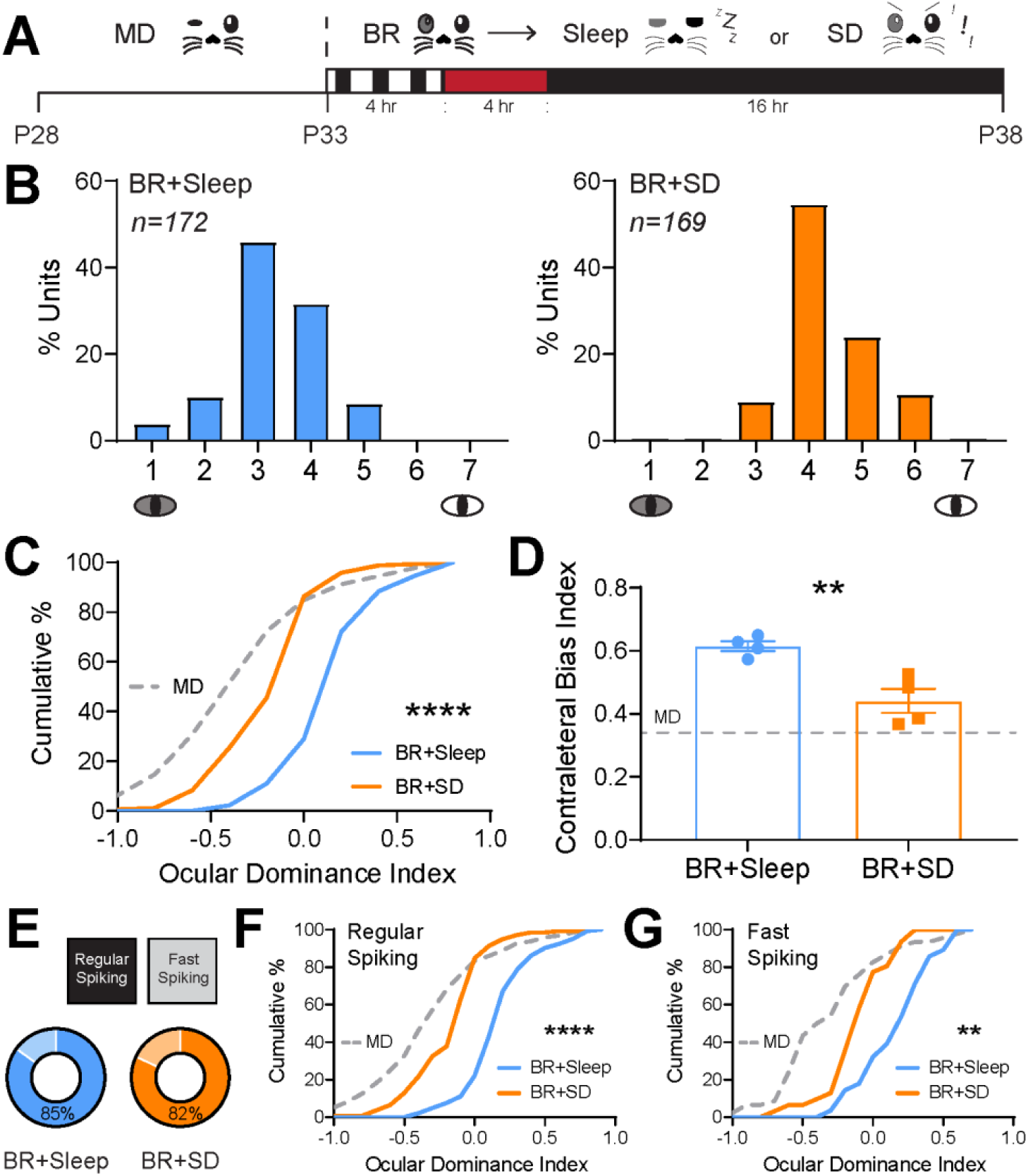
Sleep loss following BR visual experience prevents ocular dominance shifts. **(A)** Experimental design. Mice underwent 5-day MD and 5-day BR; each day after 4-hr BR, BR+Sleep mice were returned to their home cage and allowed *ad lib* sleep under infrared light, BR+SD mice underwent 4 hours of sleep deprivation through gentle handling under infrared light. **(B)** Ocular dominance histograms for bV1 neurons recorded from BR+Sleep and BR+SD groups (4 mice/group). **(C)** Cumulative distribution of ocular dominance index values for bV1 neurons recorded from BR+Sleep and BR+SD mice. **** indicates *p* < 0.0001, K-S test. Values from neurons recorded in MD-only mice from **Fig. 1** are shown (dashed gray lines) for comparison. **(D)** Contralateral bias index values were reduced for bV1 neurons recorded from BR+SD mice. Unpaired t-test: *p* = 0.0059. Error bars indicate mean ± SEM. **(E)** Proportion of recorded neurons identified as RS neurons or FS interneurons for the two groups. **(F-G)** Ocular dominance index values for recorded RS neurons **(F)** and FS interneurons **(G)** were reduced in BR+SD mice. ** and **** indicate *p* < 0.01 and *p* < 0.0001, K-S test.

### BR-mediated renormalization of DE and SE responses are reversed by sleep loss

To determine how SD affects visual responsiveness in DE and SE pathways, we assessed how visually-evoked firing rates were affected by post-BR sleep vs. SD. In both RS neurons and FS interneurons, post-BR SD led to a significant reduction of DE firing rate responses compared with those recorded from freely-sleeping mice (**Fig. 5A top** and **4D top**). When DE responses were compared across bV1 as a whole, those recorded from BR+SD mice were significantly lower than those recorded from BR+Sleep mice, similar to those recorded from mice after MD alone. Among RS neurons, we found that this effect was most pronounced in layers 2/3, where DE-driven firing rates in BR+SD mice were similar to those recorded from MD mice (**Fig. 5B**). Among FS interneurons, SD effects were most pronounced in layer 4, where depressed DE responses were similar to those of MD-only mice (**Fig. 5E**).

**Fig. 5.**
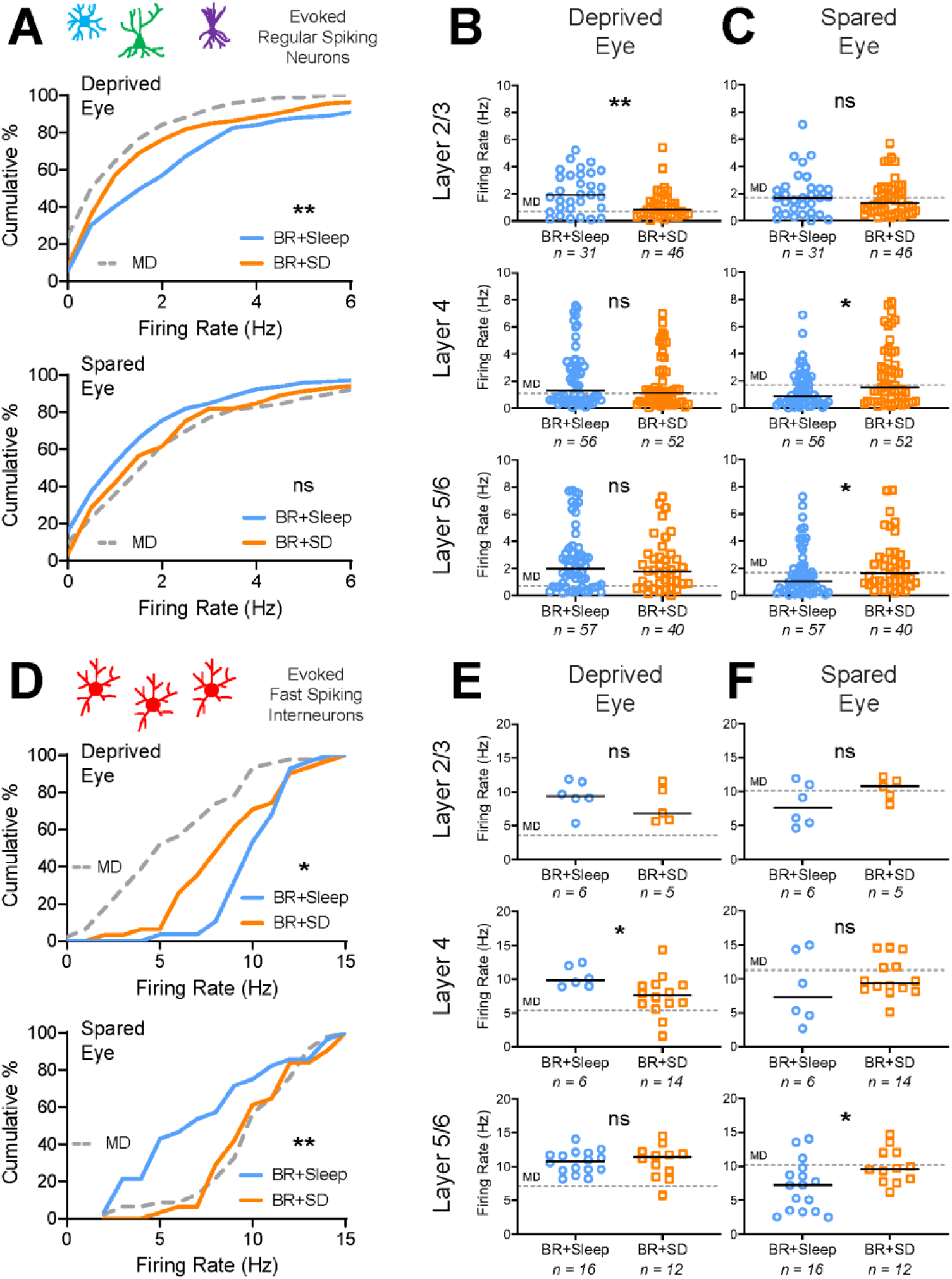
Post-BR SD prevents recovery of DE and SE responses after MD. **(A)** Cumulative distributions of maximal DE **(top)** and SE **(bottom)** visually-evoked firing rate responses for bV1 RS neurons. DE firing rate responses were significantly decreased in BR+SD mice relative to BR+Sleep mice. ** indicates *p* < 0.01, K-S test. **(B-C)** RS neurons’ DE **(B)** and SE **(C)** visually-evoked responses recorded from neurons in bV1 layers 2/3, 4, or 5/6. Solid lines indicate median values for each neuron population; * and ** indicate *p* < 0.05 and *p* < 0.01, Mann-Whitney test. **(D)** Cumulative distributions of maximal DE **(top)** and SE **(bottom)** visually-evoked firing rate responses for bV1 FS interneurons. Firing rate responses for DE and SE stimulation were significantly decreased and increased, respectively, in BR+SD mice. * and ** indicate *p* < 0.05 and *p* < 0.01, K-S test. **(E-F)** FS interneurons’ DE **(E)** and SE **(F)** visually-evoked responses recorded from neurons in bV1 layers 2/3, 4, or 5/6. Solid lines indicate median values for each neuron population. * indicates *p* < 0.05, Mann-Whitney test. Values for the MD-only condition (gray dashed lines) from **Fig. 2** are shown for comparison.

Across bV1 as a whole, RS neurons’ SE responses were not significantly different between BR+Sleep and BR+SD mice, despite a trend for higher firing rate responses after SD (**Fig. 5A bottom**). SE responses were significantly elevated in RS neurons recorded in layer 4 and layers 5/6 from BR+SD mice, where median response rates were similar to those recorded in MD-only mice (**Fig. 5C**). Across the bV1 FS interneuron population, SD interfered with BR-driven normalization of SE responses, which remained elevated, similar to those recorded from mice following MD alone (**Fig. 5D bottom**). FS interneurons’ SE responses in BR+SD mice were significantly elevated relative to BR+Sleep mice in layers 5/6, with median firing rate responses similar to those seen in MD0only mice (**Fig. 5F**). Together, these data suggest that eye-specific response renormalization due to BR in both RS neurons and FS interneurons is suppressed by post-BR SD.

To further characterize layer- and cell type-specific changes in visual responses after post-BR sleep vs. SD, we next quantified DE-driven cFos expression in bV1 of BR+Sleep and BR+SD mice, using the same DE visual stimulation strategy described for **Fig. 3**. Across bV1 as a whole, overall DE-driven cFos expression was significantly reduced in BR+SD mice compared to BR+Sleep mice (**Fig. 6A**). This reduction was most dramatic in layers 2/3 and 5/6 (**Fig. 6D**), where cFos levels in BR+SD mice were intermediate between those of BR+Sleep and MD-only mice. The density of cFos+ PV+ interneurons was likewise significantly decreased after DE stimulation in BR+SD mice (**Fig. 6C**), with dramatic reductions in layers 2/3 and 4 (**Fig. 6F**). Taken together, out data suggest that most of the changes to DE and SE responses initiated in bV1 by MD are sustained when BR is followed by SD. Conversely, BR-mediated recovery of binocular function in bV1 RS neurons and FS interneurons relies on post-BR sleep.

**Fig. 6.**
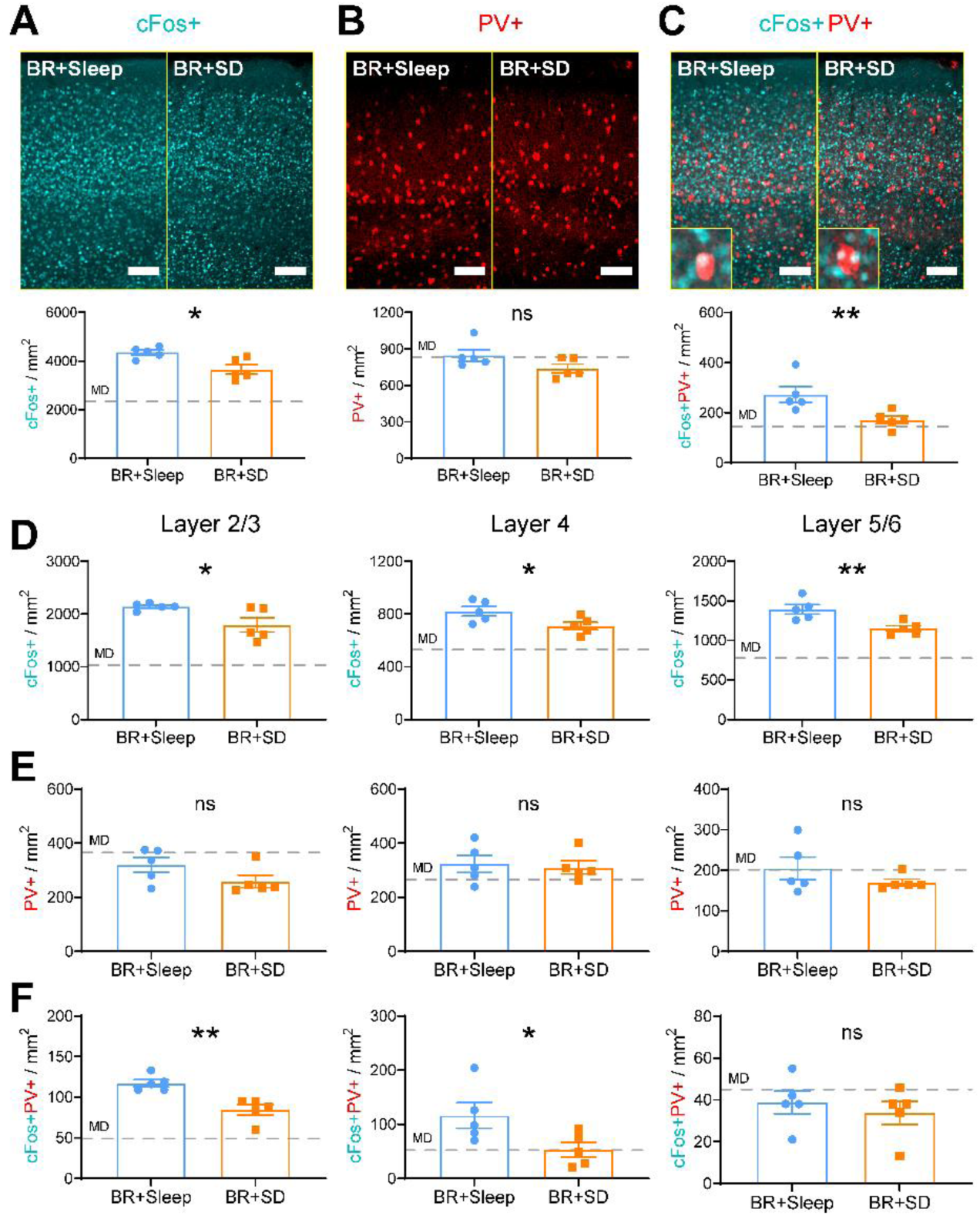
Post-BR SD prevents recovery of DE-driven cFos expression in bV1. **(A)** bV1 cFos (cyan) after DE stimulation in BR+Sleep and BR+SD mice. Mice (*n* = 5/treatment group) received DE-only visual stimulation for 30 min, then were returned to their home cages for 90 min prior to perfusion. DE-driven cFos+ neuron density was reduced in BR+SD mice relative to BR+Sleep mice. Unpaired t-test: *p* < 0.05. **(B)** PV immunostaining (red) was similar between groups. **(C)** cFos+ PV+ interneuron density was decreased in BR+SD mice relative to BR+Sleep mice. Unpaired t-test: *p* < 0.01. **(D)** cFos+ neuron density was reduced in bV1 layers 2/3, 4, and 5/6 after BR+SD relative to BR+Sleep. Unpaired t-test: *p* < 0.05, *p* < 0.05, and *p* <0.01, respectively. **(E)** PV+ interneuron density in bV1 layers 2/3, 4, and 5/6 was similar between groups. **(F)** cFos+ PV+ interneuron density was reduced in bV1 layers 2/3 and 4 in BR+SD mice relative to BR+Sleep mice. Unpaired t-test: *p* < 0.05, *p* < 0.05, and ns, respectively. Scale bar = 100 μm. Values for the MD-only condition (gray dashed lines) from **Fig. 3** are shown for comparison.

## Discussion

In this study, we compared the effects of equal-duration, qualitatively-similar binocular vs. monocular visual experience on recovery of bV1 responses following MD, using single-unit electrophysiology and immunohistochemistry. We also characterized the effects of post-experience sleep and sleep loss on these processes. Critically, a side-by-side comparison of equal-duration BR and RO clearly showed that bV1 ocular dominance shifts in favor of the SE are reversed after 5-day BR, but not 5-day RO (**Fig. 1**). This reversal is present in both RS neurons and FS interneurons, and is associated with reversal of both MD-driven DE response depression and SE response potentiation (**Fig. 2, Fig. 3**). Insofar as MD serves as a model for amblyopia caused by disruption of vision in one of the two eyes during childhood, these data add to a body of growing evidence that suggests that binocular visual experience may offer advantages over patching of the dominant eye, which until recently was the standard of care for amblyopia. However, when daily BR experience was followed by SD, recovery of normal binocular vision in bV1 after MD was nearly completely blocked (**Fig. 4-6**). This suggests that the relative timing of sleep relative to recovery experience is potentially a critical – but overlooked - consideration for the treatment of amblyopia.

How do BR and RO differ in their effects on the bV1 circuit? Here we find that 5-day MD causes significant DE response depression among both RS neurons and FS interneurons in bV1 – with the most dramatic depression observed in layers 2/3. These changes not only reduce firing rates, but also strongly reduce DE-driven cFos expression among PV+ and PV-neuron populations in layers 2/3. These findings are consistent with results of longitudinal calcium imaging studies in mouse V1 (8) and longitudinal electrophysiological recordings in cat V1 (7). While this response depression is completely reversed by 5-day BR, only partial recovery of DE responses is achieved with RO. Differential recovery between BR and RO is evident both at the level of firing rates (**Fig. 2**) and DE-driven cFos expression (**Fig. 3**). Across the initial 5-day MD period, both FS interneurons and RS neurons also show widespread potentiation of SE responses, across all layers of bV1. These SE response changes (which are thought to occur only after DE response depression has already take place) (5, 7, 36) appear to be almost fully reversed after 5-day BR. In contrast, SE response enhancement is minimally altered after 5-day subsequent RO. In general, these findings are consistent with intrinsic signal imaging studies in binocular mouse V1, which indicated that a single day of BR is superior to RO at restoring binocularity (29). Future studies will be needed to determine the extent to which initial eye-specific response changes during MD are mediated by Hebbian vs. homeostatic plasticity mechanisms (46-52), and how differential outcomes with BR vs. RO themselves reflect Hebbian vs. homeostatic changes within bV1.

How does post-experience sleep or sleep loss affect bV1 during recovery? Our data clearly demonstrate that following periods of BR experience, subsequent sleep is essential for full recovery of MD-driven changes in ocular dominance (**Fig. 4**), DE and SE firing rate responses (**Fig. 5**), and DE-driven cFos expression (**Fig. 6**). These are the first data demonstrating that following MD, sleep plays a critical role in restoring normal visual function. Prior work has shown that post-MD sleep is essential for initial ocular dominance shifts in favor of the spared eye (7, 35-38) and for MD-induced structural plasticity in V1 neurons (45). However, the role of sleep in promoting recovery of function following the onset of amblyopia is understudied. Prior work done in cats after brief RO indicated that post-RO SD had little impact – and even tended to reduce recovery of binocular vision (39). Less is known about interactions between BR visual experience and subsequent sleep. While post-BR sleep has been suggested to promote homeostatic downscaling of firing rates in rodent monocular zone (53), no prior work has addressed how it affects bV1 ODP. The present work characterizes how sleep contributes to experience-driven recovery of binocular vision in bV1. We find that post-BR sleep is required for reversal of both DE response depression and SE response potentiation, in both RS neurons and FS interneurons. As with changes driven by initial MD (**Fig. 2-3**), changes driven by post-BR sleep appear to be most dramatic in layers 2/3 (**Fig. 5-6**).

Why might sleep be essential for these changes? Available data suggests that both Hebbian synaptic potentiation and weakening can occur in bV1 during post-MD sleep (7, 36, 37, 54, 55) through sleep-dependent activation of specific molecular pathways (36, 37, 56) or sleep-specific activity patterns (7). It is plausible that similar mechanisms are involved in the reversal of MD-driven synaptic changes during post-BR sleep. For example, specific oscillatory patterning of neuronal firing in the V1-LGN network during sleep may be essential for spike timing-dependent plasticity between synaptically-connected neurons (33, 34, 57-60).

Alternatively, sleep may promote permissive changes in biosynthetic pathways that are essential for consolidating some forms of plasticity *in vivo* (61-63). In V1, sleep plays a role in increasing inhibition within layers 2/3, reducing E/I ratios across the rest phase (64); this may play a role in reversing ocular dominance changes driven by suppression of FS interneurons in the context of MD (7, 8). Finally, sleep also contributes to homeostatic changes in V1 neurons’ firing rates (53, 65); thus sleep-dependent homeostatic plasticity may also contribute to bV1 changes observed in BR+Sleep, but not BR+SD, mice.

Many factors affect the degree of ODP initiated by MD in animal models of amblyopia, including behavioral state (35, 36, 38) and neuropharmacology (10, 66-68). Emerging data suggests that these factors may also affect recovery from amblyopia (67, 69, 70). However, numerous findings have raised debate about whether dominant-eye (SE) patching, the decades-long standard of care for amblyopia, provides the optimal sensory stimulus for promoting recovery of vision (18, 22, 71, 72). Here, in side-by-side comparison of the effects of brief binocular vs. monocular recovery experience, we show the two have strikingly different effects on bV1 ocular dominance. Critically, our data also provide the first demonstration that the timing of sleep relative to visual experience during amblyopia treatment may be critical for restoring normal bV1 function. We hope that these data will inform future strategies for optimizing amblyopia treatment in children, to lessen the long-term impact of early-life visual disruptions.

## Materials and Methods

### Animal housing and husbandry

All mouse husbandry and experimental/surgical procedures were reviewed and approved by the University of Michigan Internal Animal Care and Use Committee. C57BL6/J mice were housed in a vivarium under 12h:12h light/dark cycles (lights on at 9AM) unless otherwise noted, and had *ad lib* access to food and water. After eyelid suture surgeries, mice were single housed in standard cages with beneficial environmental enrichment. For studies comparing the effects of sleep on BR visual experience, mice were housed with a 4h:20h light:dark cycle (lights on from 9AM-1PM during visual enrichment, dim red light outside of visual enrichment) and had *ad lib* access to food and water.

### Monocular deprivation, recovery, visual enrichment, and sleep deprivation

For monocular deprivation (MD), mice were anesthetized at P28 using 1-1.5% isoflurane. Nylon non-absorbable sutures (Henry Schien) were used to occlude the left eye. Sutures were checked twice daily to verify continuous MD; during this time they were handled 5 min/day. After MD (at P33), mice were anesthetized with 1-1.5% isoflurane a second time and left eyelid sutures were removed. Mice that underwent binocular recovery (BR) were then housed over the next 5 days with both eyes open; during this time they were handled daily for 5 min/day. Mice that underwent reverse occlusion (RO) had the right (previously spared; SE) eye sutured for the next 5 days; these mice were also handled 5 min/day during this period. Mice that lost sutures during the MD or recovery periods or developed eye abnormalities were excluded from the study. BR and RO mice underwent a similar 5-day period of daily enriched visual experience from P34-38. This regimen consisted of a daily placement in a 15” × 15” Plexiglas chamber surrounded by 4 high-contrast LED monitors, from ZT0 (lights on) to ZT4. Phase-reversing oriented grating stimuli (0.05 cycles/deg, 100% contrast, 1 Hz reversal frequency) of 8 orientations were presented repeatedly on the 4 monitors in a random, interleaved fashion. During this 4-h period of daily visual enrichment, mice were encouraged to remain awake and explore the chamber via presentation of a variety of enrichment toys (novel objects, transparent tubes, and a running wheel) and palatable treats. For SD studies on BR experience, following the 4-h visual enrichment period, mice were placed in their home cage under dim red light (to prevent additional visual experience), and were sleep deprived (SD) by gentle handling (57, 65). Briefly, gentle handling procedures involved visually monitoring the mice for assumption of sleep posture - i.e., huddled in their nest with closed eyes. Upon detection of sleep posture, the cage was either tapped or (if necessary) shaken briefly (1-2 seconds). If sleep posture was maintained after these interventions, the nesting material within the cage would be moved using a cotton-tipped applicator. No novel objects or additional sensory stimuli were provided, to limit sensory-based neocortical plasticity during sleep deprivation procedures (73). As previously described, similar procedures used for sleep deprivation in adult mice (74) and critical period cats (38) either have no significant effect on serum cortisol, or increases it to a degree that is orders of magnitude lower than that capable of disrupting ODP (75).

### *In vivo* neurophysiology and single unit analysis

Mice underwent stereotaxic, anesthetized recordings using a combination of 0.5-1.0% isoflurane and 1 mg/kg chlorprothixene (Sigma). A small craniotomy (1 mm in diameter) was made over right-hemisphere bV1 (i.e., contralateral to the original DE) using stereotaxic coordinates 2.6-3.0 mm lateral to lambda. Recordings of neuronal firing responses were made using a 2-shank, linear silicon probe spanning the layers of bV1 (250 μm spacing between shanks, 32 electrodes/shank, 25 μm inter-electrode spacing; Cambridge Neurotech). The probe was slowly advanced into bV1 until stable, consistent spike waveforms were observed on multiple electrodes. Neural data acquisition using a 64-channel Omniplex recording system (Plexon) was carried out for individual mice across presentation of visual stimuli to each of the eyes, via a full field, high-contrast LED monitor positioned directly in front of the mouse. Recordings were made for the right and left eyes during randomly interleaved presentation of a series of phase-reversing oriented gratings (8 orientations + blank screen for quantifying spontaneous firing rates, reversing at 1 Hz, 0.05 cycles/degree, 100% contrast, 10 sec/stimulus; Matlab Psychtoolbox). Spike data for individual neurons was discriminated offline using previously-described PCA and MANOVA analysis (57, 65, 76, 77) using Offline Sorter software (Plexon).

For each visually-responsive neuron, a number of response parameters were calculated (7, 36). An ocular dominance index was calculated for each unit as (C-I)/(C+I) where C represents the maximal visually-evoked firing rate for preferred-orientation stimuli presented to the contralateral (left/deprived) eye and I represents the maximal firing rate for stimuli presented to the ipsilateral (right/spared) eye. Ocular dominance index values range from −1 to +1, where negative values indicate an ipsilateral (SE) bias, positive values indicate a contralateral (DE) bias, and values close to 0 indicate similar responses for stimuli presented to either eye.

### Histology and immunohistochemistry

Following all electrophysiological recordings, mice were euthanized and perfused with ice cold PBS and 4% paraformaldehyde. Brains were dissected, post-fixed, cryoprotected in 30% sucrose solution, and frozen for sectioning. 50 μm coronal sections containing bV1 were stained with DAPI (Fluoromount-G; Southern Biotech) to verify electrode probe placement in bV1. Mice whose electrode placement could not be verified were excluded from further analyses.

For immunohistochemical quantification of PV and DE-driven cFos expression in bV1, mice from all groups underwent monocular eyelid suture of the original SE at ZT12 (lights off) the evening before visual stimulation. At ZT0 (next day), DE stimulation was carried out in the LED-monitor-surrounded arena with treats and toys to maintain a high level of arousal (as described above) Mice from all groups were exposed to a 30-min period of oriented gratings (as described for visual enrichment above), after which they were returned to their home cage for 90 min (for maximal visually-driven cFos expression) prior to perfusion. Coronal sections of bV1 were collected as described above and immunostained using rabbit-anti-cFos (1:1000; Abcam, ab190289) and mouse-anti-PV (1:2000; Millipore, MAB1572) followed by secondary antibodies: Alexa Fluor 488 (1:200; Invitrogen, A11032) and Alexa Fluor 594 (1:200; Invitrogen, A11034). Stained sections were mounted using Prolong Gold antifade reagent (Invitrogen) and imaged using a Leica SP8 confocal microscope with a 10× objective, to obtain z-stack images (10 μm steps) for maximum projection of fluorescence signals. Identical image acquisition settings (e.g. exposure times, frame average, pixel size) were used for all sections. cFos+ and PV+ cell bodies were quantified in 3-4 sections (spanning the anterior-posterior extent of bV1) per mouse by a scorer blinded to animal condition and reported as approximate density, using previously established procedures. Co-labeling was quantified using the Image J JACoP plugin (78) and values for each mouse are averaged across 3-4 sections.

### Statistical analysis

Statistical analyses were carried out using GraphPad Prism software (Version 9.1). Comparisons of ocular dominance, firing rates, and visual response properties were made using stably recorded, visually-responsive units in bV1. Nonparametric tests were used for non-normal data distributions. Specific statistical tests and p-values can be found within the results section and in corresponding figures and figure legends.

## Acknowledgments

We thank Abbey Roelofs (LSA Technology Services) for software programming assistance. This work was supported by a University of Michigan Candidate Grant and a Rackham Merit Fellowship to J.D.M., NIH R01NS104776, and a Research to Prevent Blindness Walt and Lilly Disney Award for Amblyopia Research to S.J.A.

